# MoBPSweb: A web-based framework to simulate and compare breeding programs

**DOI:** 10.1101/2020.07.08.193227

**Authors:** T. Pook, L. Büttgen, A. Ganesan, N.T. Ha, H. Simianer

## Abstract

Selective breeding is a continued element of both crop and livestock breeding since early prehistory. In this work, we are proposing a new web-based simulation framework (“MoBPSweb”) that is combining a unified language to describe breeding programs with the simulation software MoBPS, standing for ‘Modular Breeding Program Simulator’. Thereby, MoBPSweb is providing a flexible environment to enter, simulate, evaluate and compare breeding programs. Inputs can be provided via modules ranging from a Vis.js-based flash environment for “drawing” the breeding program to a variety of modules to provide phenotype information, economic parameters and other relevant information. Similarly, results of the simulation study can be extracted and compared to other scenarios via output modules (e.g. observed phenotypes, accuracy of breeding value estimation, inbreeding rates). Usability of the framework is showcased along a toy example of a dairy cattle breeding program on farm level, with comparing scenarios differing in implemented breeding value estimation, selection index and selection intensity being considered. Comparisons are made considering both short and long-term effects of the different scenarios in terms of genomic gains, rates of inbreeding and the accuracy of the breeding value estimation. Lastly, general applicability of the MoBPSweb framework and the general potential for simulation studies for genetics and in particular in breeding are discussed.

## Introduction

**S**ince early prehistory, selective breeding has been a tool that humanity has used to, among others, ensure the food supply with examples spanning back to the development from teosinte to maize in Mesoamerica and animal breeding for war horses and dogs in the Roman Empire (Sidnell 2007). Over time, breeding has become more refined with concepts like genetic inheritance (Mendel 1866), quantitative genetics (Galton 1889; Fisher 1918) and population genetics (Wright 1922) being introduced. Today, breeders have a large toolbox of breeding technologies, ranging from high-throughput genotyping and phenotyping (Solberg *et al*. 2006; Cabrera-Bosquet *et al*. 2012) to highly advanced bio technology (Jinek *et al*. 2012) to complex quantitative models for breeding value estimation and QTL detection (Meuwissen *et al*. 2001; Klein *et al*. 2005; VanRaden 2008). These advanced methods are both a blessing and a curse, as managing and optimizing such a breeding program is a highly complex problem (Henryon *et al*. 2014).

In recent years a variety of tools to simulate breeding programs (Sargolzaei and Schenkel 2009; Faux *et al*. 2016; Liu *et al*. 2019; Pérez-Enciso *et al*. 2020; Pook *et al*. 2020) have been developed to aid breeders in their decision as they provide a controlled and repeatable environment to modify individual parameters of a breeding scheme and by that draw conclusion on their impact on the breeding objective. Potential goals usually include trait complexes such as productivity, fitness, adaptation and inbreeding.

A common problem of simulation programs is that they either are not flexible enough to simulate complex breeding programs or that the extremely high number of necessary options makes the use of such a software difficult (e.g. the main function of the software MoBPS (Pook *et al*. 2020) *breeding*.*diploid()* has over 200 parameters). Therefore, simulation studies are often times carried out via new and self-written code that is tailor-made for the problem at hand. However, this leads to the problem of inefficient and error prone code that potentially neglects less-intuitive but still important factors that nonetheless are impacting the breeding scheme.

In a companion paper, Simianer *et al*. (2020) proposed a unifying concept to describe breeding programs. Even though the focus in that work was mostly on animal breeding, concepts can readily be extended to plant breeding. Key idea of the concept is to describe each breeding program via a set of nodes and edges, with nodes representing cohorts of individuals and edges representing a set of potential breeding actions that can be taken in breeding. In the following, we will use the notation as proposed in Simianer *et al*. (2020).

In this work, we combine the ideas of Simianer *et al*. (2020) with the highly efficient breeding program simulator MoBPS (Pook *et al*. 2020) to provide a web-based application (“MoBP-Sweb”) to enter, simulate, evaluate and compare breeding programs in an user-friendly and intuitive environment.

## Material and Methods

MoBPSweb is a web-based application running on a NodeJS server that uses the VueJS and Express frameworks in the fronend and backend, respectively. We further employ a MongoDB as a backend database (Chodorow 2013). The simulation of the breeding program and all downstream analysis are performed in R (R Core Team 2017) by using wrapper functions of the MoBPS R-package. An overview of the modules contained in the framework is given in Figure 1 with all sub-modules being described in more detailed in the following subsections.

**Figure 1.**
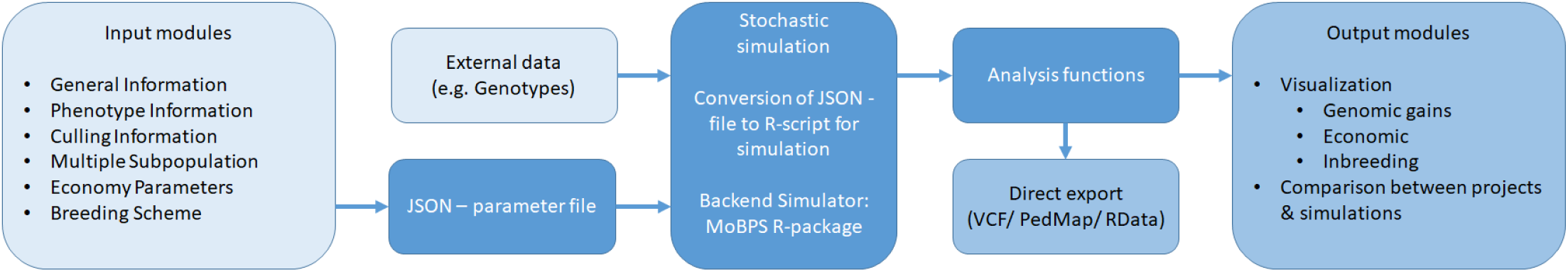
Schematic overview of the MoBPSweb framework.

### Breeding scheme

As described in Simianer *et al*. (2020) a breeding scheme can be represented by a set of cohorts (nodes) and breeding actions (edges). This is implemented in an interactive environment based on Vis.js. To detect potentially issues (e.g. loops, missing required information), the resulting set of nodes and edges is constantly checked and potential issues are reported as warnings.

### Input modules

In addition to the breeding scheme itself, we provide a variety of different input modules to enter further information:

- General Information
- Phenotype Information
- Culling Information
- Multiple Subpopulations
- Economy Parameters

With the exception of the “General Information”-module all of these modules are optional. The basic functionality of each respective input modules is described in the following subsections.

### General Information

In the “General Information”-module basic information like the species and the underlying genomic map (including used arrays) are chosen. Maps can either be uploaded, manually created or imported from the Ensembl database (Zerbino *et al*. 2018).

### Phenotype Information

The “Phenotype Information”-module provides options to generate a set of traits. This includes defining phenotypic mean and variance, as well as the heritability, repeatability, number of underlying QTL for each trait. Potential correlations between traits need to be given for both the genetic and residual component. Furthermore, selection indices to later perform selection based on multiple traits can be defined here. Exemplary sets of traits for a variety of species are provided as templates.

### Culling Information

The “Culling Information”-module is designed to model individuals leaving the breeding nucleus. In contrast to breeding actions via edges, no new individuals are generated. Input parameters are in line with parameters of the culling options in *breeding*.*diploid()* in the MoBPS R-package with the added benefit that the age of individuals is tracked and culling actions are automatically performed as soon as an individual reaches a given age.

### Multiple Subpopulations

The “Multiple Subpopulation”-module is designed to generate founder individuals of different genetic origin, e.g. for representing different breeds or lines in a crossbreeding scenario. This is only relevant in case no genotype information is imported for some of the founders and will lead to simulated genotypes being drawn from different per marker allele frequencies.

### Economy Parameters

In the “Economy Parameters”-module basic information regarding the cost of different breeding actions can be entered. This includes both fix costs and variable costs like genotyping, phenotyping and housing costs that are automatically discounted according to provided interest rates.

### JSON to R conversion

Output of the web-interface is a JavaScript Object Notation (JSON)-file containing all entered information of the breeding scheme. In the following this JSON-file needs to be translated into interpretable R-code. All this is implemented in the function *json*.*simulation()* in the R-package MoBPS (Pook *et al*. 2020). The translation procedure for all advanced modules is relatively straight-forward, as most parameters directly correspond to a parameter in the MoBPS R-package. E.g. the size of the genome and the selected underlying array will set the map parameter in *creating*.*diploid()* and the phenotyping module is feeding in information for a separate call of *creating*.*trait()* from the MoBPS R-package.

The conversion of the breeding scheme itself is done by first detecting if the breeding scheme has any repeat edges. If that is the case, it will subsequently check which nodes can be generated without use of any repeat. Next, all repeats that can be executed based on the already available nodes are executed by generating copies of all nodes between the node of origin and the target node of the repeat (including the node of origin and excluding the target node). Nodes generated via repeat are serial numbered via “_1”, “_2” etc. to indicate the repeat number. This procedure is repeated until all repeat edges are resolved, leading to a breeding program without any repeats remaining.

Next, the actual breeding scheme is simulated by first generating all founder nodes via *creating*.*diploid()*. All remaining cohorts are generated by separate calls of *breeding*.*diploid()*. All necessary information for this is stored in incoming edges and the node itself. For edges this includes the breeding type, but also respective subsequent details (e.g. for selection: method for breeding value estimation, cohorts used for breeding value estimation etc.). For nodes, information includes the phenotyping class, the share of genotyped individuals and the housing class.

The order of generation will be derived based on generation times assigned to each edge and cross-dependencies (e.g. when phenotyping information from one cohorts is needed to generate breeding values for another cohort).

### Evaluating simulation outputs

After successfully simulating a breeding program, the resulting population list is stored and can be analysed. In the web-interface itself, we are providing five implemented analysis modules that are each corresponding to an analysis function from the MoBPS R-package:

- Observed Phenotypes - *get*.*pheno()*
- True Breeding Values - *get*.*bv()*
- Accuracy of Breeding Value Estimation - *analyze*.*bv()*
- Relationship and Inbreeding within Cohorts - *kinship*.*emp*.*fast()*
- Major QTLs - *get*.*geno()*

All these modules can either be applied on a single run of the simulation or be averaged across multiple random replicates of the same simulation scenario. Furthermore, the “CompareProject”-module is providing an environment to compare different breeding schemes with each other by use of the same analysis functions.

Note that it is also possible to directly extract pedigree and phenotype information, VCF (Danecek *et al*. 2011) / PedMap (Purcell *et al*. 2007)-files for selected cohorts and the population list itself, and to proceed with own analyses in R or similar programs.

### Data Availability

All code underlying MoBPSweb can be found at https://github.com/tpook92/MoBPS_web. A running version of MoBPSweb can be found at www.mobps.de. JSON-files for all compared scenarios discussed can be found in Supplementary File S1 with the reference also being included as a template at www.mobps.de. Figure S1-S4 display the genomic gain for the traits RZR, RZE, RZS, RZKm for the reference dairy cattle breeding scheme. Supplementary files are available at FigShare.

## Results

### Base-line scenario

In the following, we will discuss an exemplary use case of the web-interface for a dairy cattle breeding program on farm level. The breeding scheme, as entered in the flash application of MoBPSweb, is given in Figure 2.

**Figure 2.**
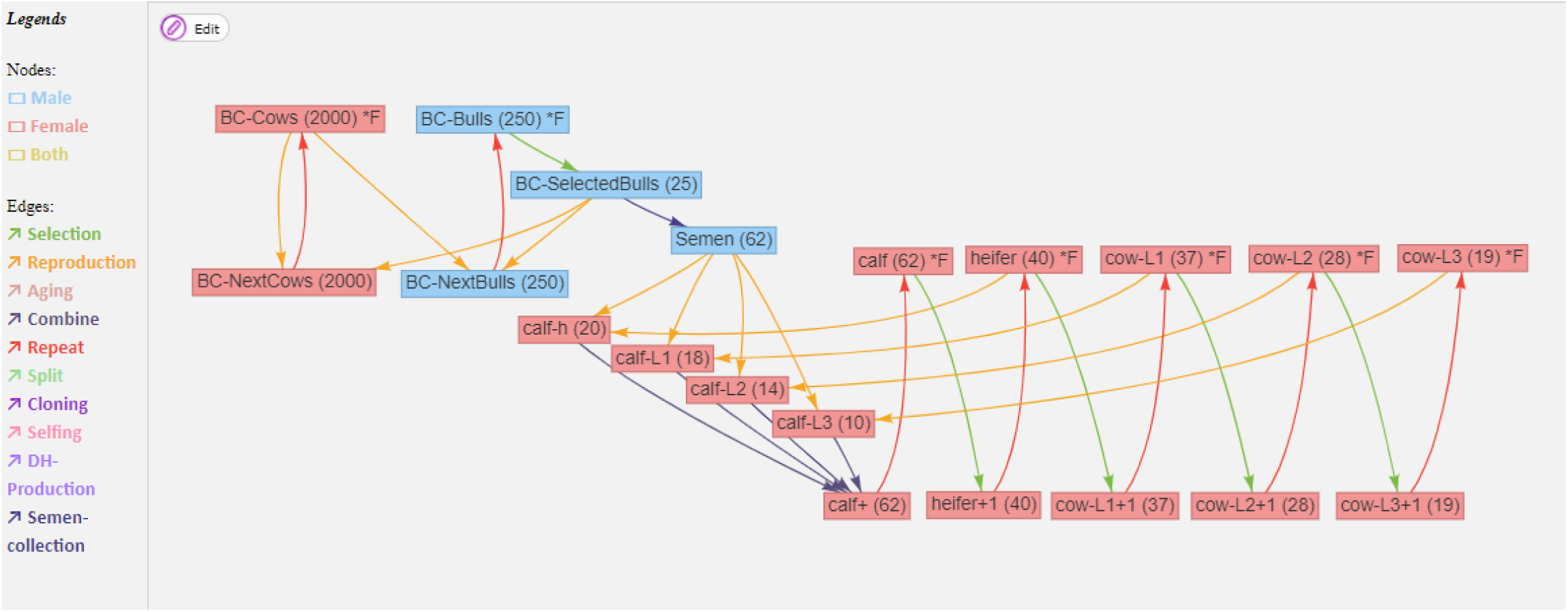
A dairy cattle selection program within a herd, with animals separated in age groups. Details on the attributes of all nodes and edges can be found in Supplementary Material S1.

At each time point the farm has spots for 184 animals consisting of five cohorts (calf, heifer, cow-L1, cow-L2, cow-L3) that are split based on age. New animals are generated by the use of semen from a breeding company with heifers and older cows being used for reproduction. These offspring are then merged into a joined cohort and subsequently used as the calf-cohort (“calf+”; Figure 2) in the next repeat. New animals for the other four cohorts are chosen by selecting the required number of animals from the respectively one year younger cohort (“heifer+1”, “cow-L1+1”, “cow-L2+1”, “cow-L3+1”; Figure 2). For this selection procedure five traits are simulated and animals are selected based on an index selection formed from those traits (Table 1) with phenotypes only partially available based on age. It is assumed that already combined traits, as being used in the standard German breeding evaluation system can be observed on each cow. Five traits (RZM, RZE, RZR, RZS, RZKm) were chosen according to Vereinigte Informationssysteme Tierhaltung w.V. (2020) and standardized to have a starting mean of 100 with a genetic standard deviation (gSD) of 12 to allow a better comparative assessment. Genetic correlations were taken from Vereinigte Informationssysteme Tierhaltung w.V. (2020) and residual correlations assumed to be the same (Table 2). Longevity traits are ignored and removed from the index as phenotyping of those traits can be far more difficult, thereby going beyond the scope of the toy example. Since longevity (RZN) with an index weight of 20 was removed from the analysis, the index weights of the remaining traits add up to 80. As traits given here already represent combinations of other traits, no heritability or repeatability values are given in Vereinigte Informationssysteme Tierhaltung (2020). Instead reasonable values were estimated based on the given sub-trait heritablities and using estimates in the literature (Roman *et al*. 2000; Oyama *et al*. 2002).

**Table 1.** Overview of simulated traits for heritability, repeatability, index weights and if traits are phenotyped for each cohort (Vereinigte Informationssysteme Tierhaltung w.V. 2020). Numbers in brackets indicate the accumulated number of observations for the respective trait.

| Trait | heritability | repeatability | Index Weighting | calf | heifer | cow-L1 | cow-L2 | cow-L3 |
| --- | --- | --- | --- | --- | --- | --- | --- | --- |
| RZM (Milk) | 0.30 | 0.50 | 45 | No | No | Yes (1) | Yes (2) | Yes (3) |
| RZE (type) | 0.25 | 0.25 | 15 | No | No | Yes (1) | Yes (1) | Yes (1) |
| RZR (fertility) | 0.02 | 0.06 | 10 | No | Yes (1) | Yes (2) | Yes (3) | Yes (4) |
| RZS (somatic cell score) | 0.15 | 0.22 | 7 | No | No | Yes (1) | Yes (2) | Yes (3) |
| RZKm (calving traits) | 0.05 | 0.09 | 3 | No | No | Yes (1) | Yes (2) | Yes (3) |

**Table 2.**
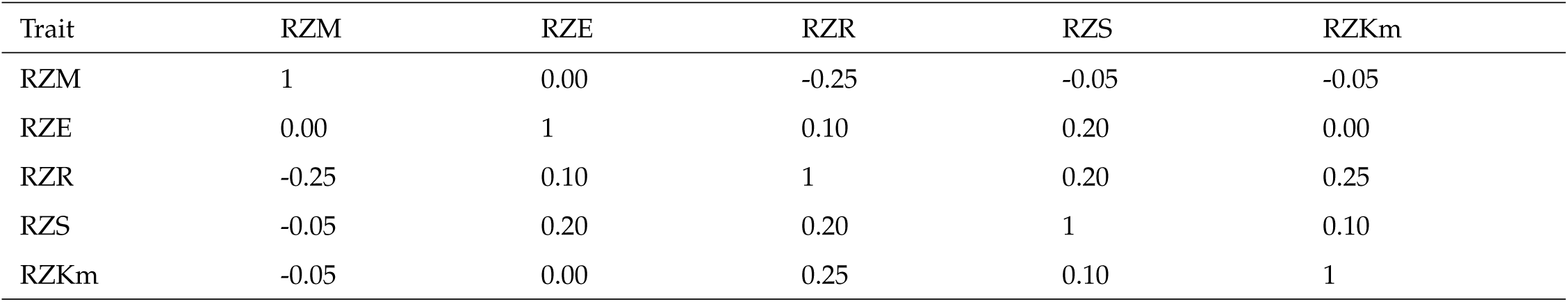
Genetic / residual correlation for considered traits given in Table 1 in the lower / upper triangle matrix.

| Trait | RZM | RZE | RZR | RZS | RZKm |
| --- | --- | --- | --- | --- | --- |
| RZM | 1 | 0.00 | -0.25 | -0.05 | -0.05 |
| RZE | 0.00 | 1 | 0.10 | 0.20 | 0.00 |
| RZR | -0.25 | 0.10 | 1 | 0.20 | 0.25 |
| RZS | -0.05 | 0.20 | 0.20 | 1 | 0.10 |
| RZKm | -0.05 | 0.00 | 0.25 | 0.10 | 1 |

In the baseline scenario, selection on the farm is assumed to be solely based on individual phenotypes and paternal breeding material is received in the form of semen from the breeding company. The underlying breeding scheme from the side of the breeding company is simplified with just one male and one female cohort per repeat and small animal numbers (250 bulls, 2000 cows) as it is not the main focus of this study. Selection for the bulls here is done via single-step breeding value estimation (Legarra *et al*. 2014) with only bulls being genotyped and phenotypes of the last two repeats of cows being used. Cows of the breeding company are assumed to have the same number of phenotypic observations per trait as cows after the third lactation period (“cow-L3”, Table 1). The breeding cycle was repeated 20 times with additional five burn-in repeats to build up some linkage disequilibrium and obtain an initial pedigree structure. An exemplary node and edges from the web interface are given in Figure 3, with details on all edges and nodes given in Supplementary File S1.

**Figure 3.**
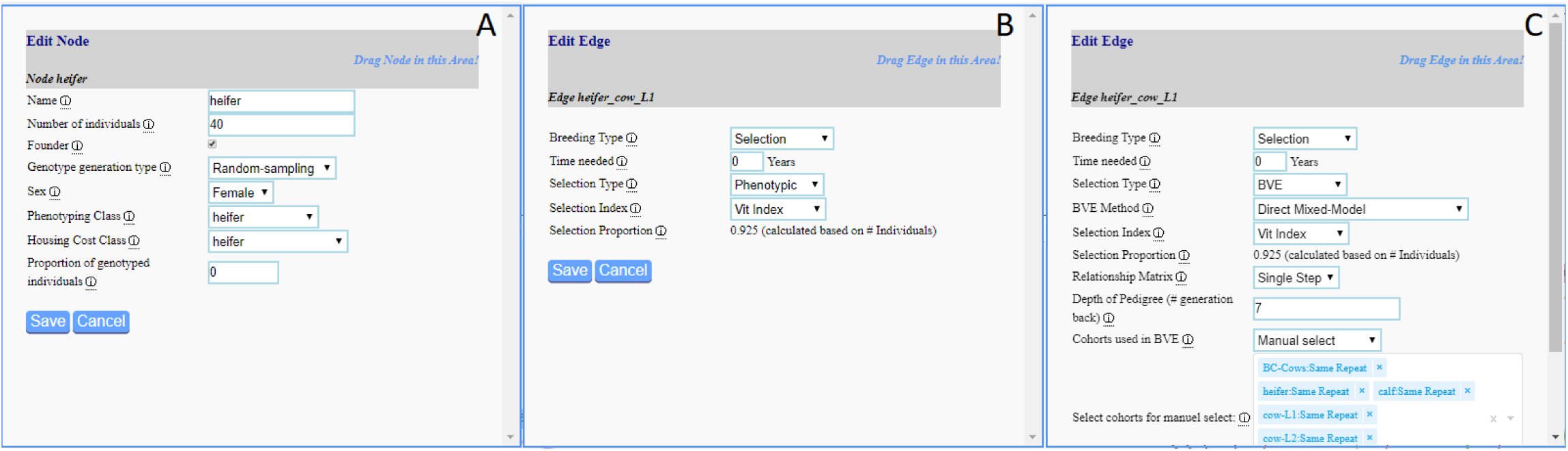
Exemplary node (A) and edge (B) of the base-line scenario (Figure 2) and genomic selection performed in scenario “ss-BLUP_BVE” (C).

As one would expect, underlying true genomic values of individuals are going up over time with by far the highest increase for RZM (10.5 gSD in 20 repeats) as the highest index weight was put on it and heritablity was also high (Figure 4). The corresponding results for the other traits are given in Supplementary Figure S1-S4. In addition, inbreeding rates are increasing by about 0.01 per cycle (Figure 5). Prediction accuracies for the breeding value estimation of the bulls is relatively low with values ranging between 0.3 for RZR and 0.6 for RZM (which could easily be increased by enlarging the training population).

**Figure 4.**
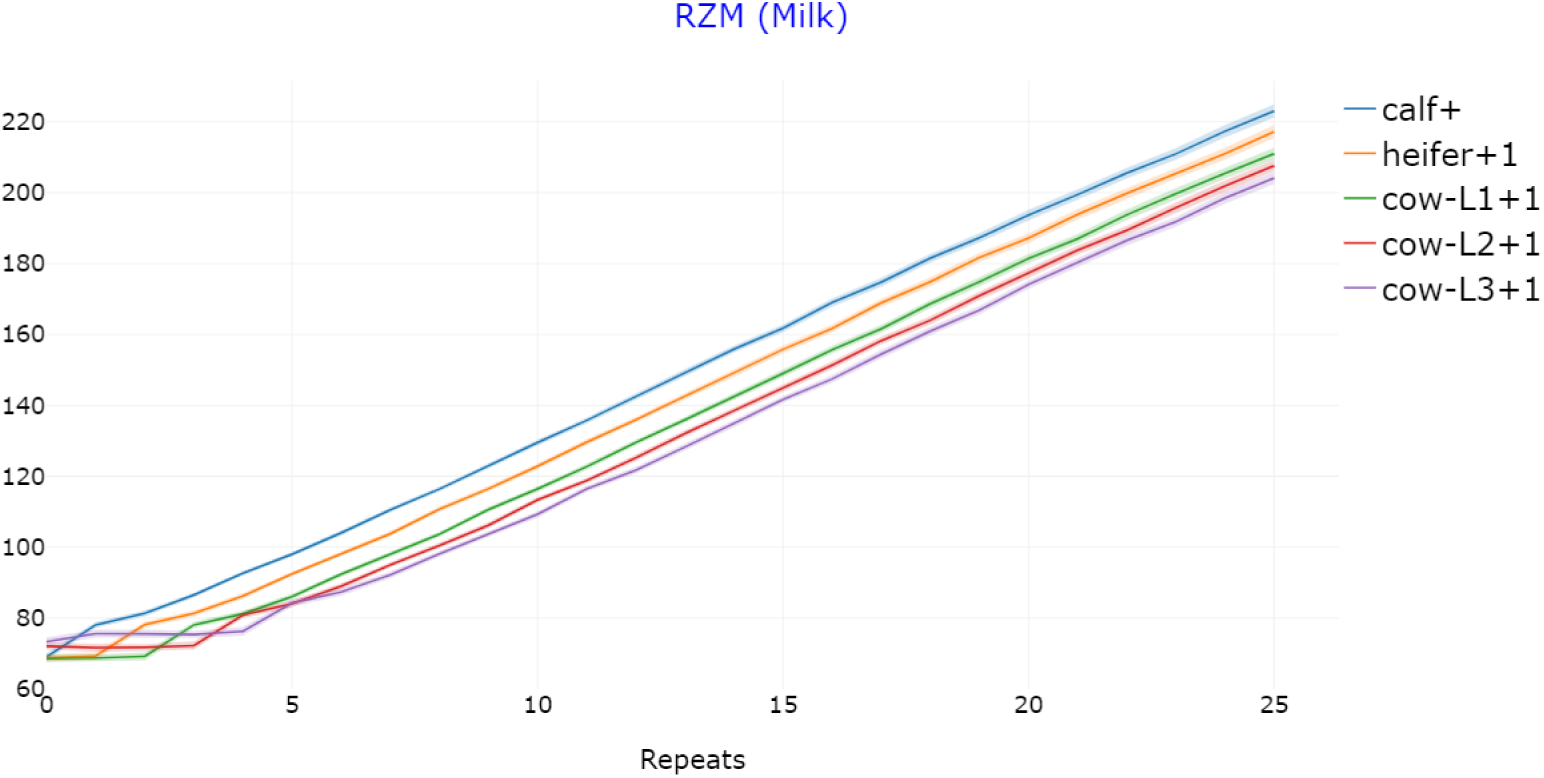
Genomic gain in the reference dairy cattle breeding scheme (Figure 2) with 95% confidence bands for the trait RZM with average genomic value standardized to 100 after 5 repeats. This figure is an exemplary output of the “True Breeding Values”-module in MoBPSweb (www.mobps.de).

**Figure 5.**
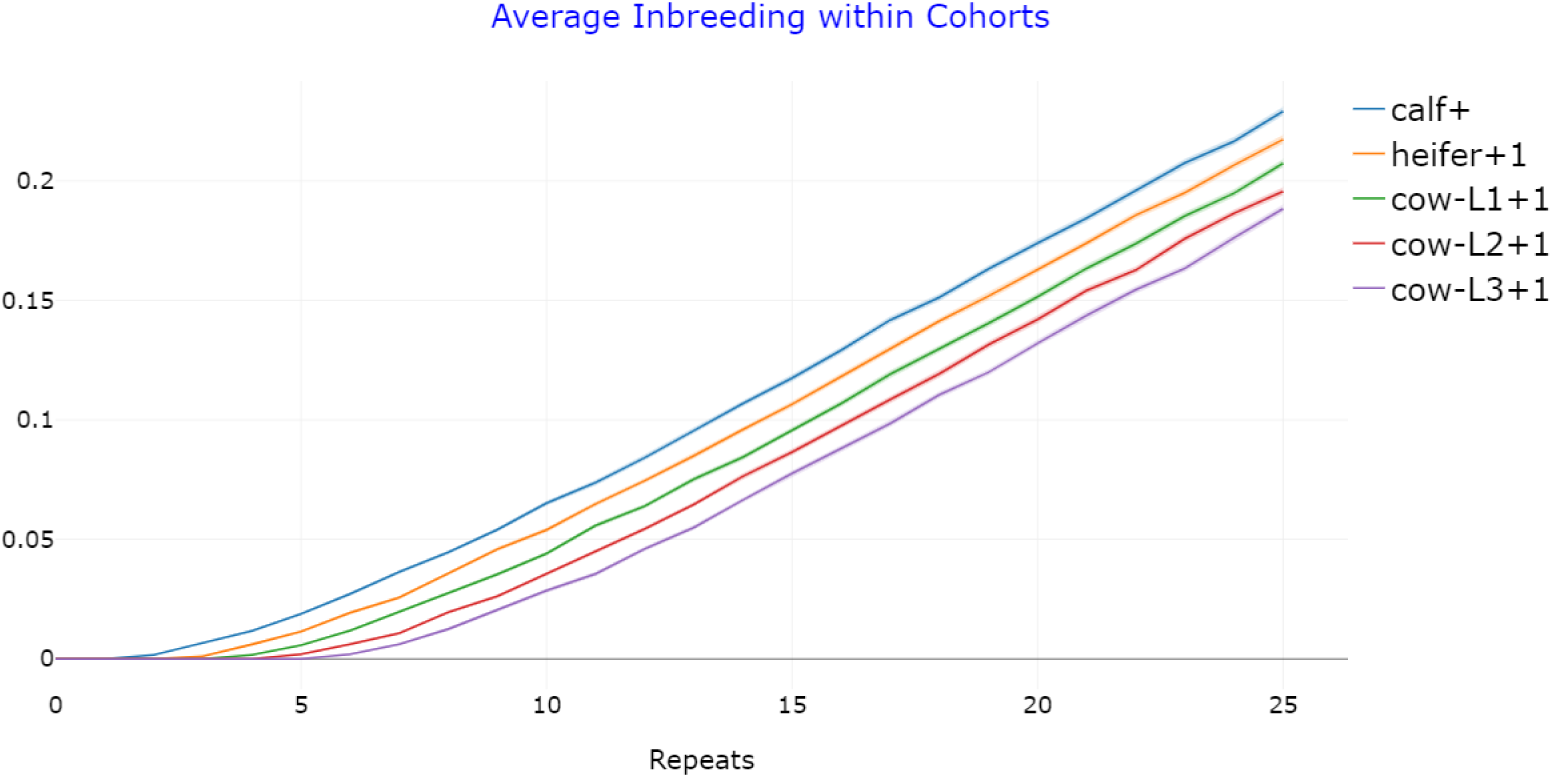
Increase in inbreeding in the reference dairy cattle breeding scheme for the calf+ cohort (Figure 2) with 95% confidence bands. This figure is an exemplary output of the “Relationship and Inbreeding within Cohorts”-module in MoBPSweb (www.mobps.de).

### Comparative scenarios

In addition to the base-line scenario, we will consider a variety of modifications of the original breeding program and analyze their impact:

1. Use of pedigree-based breeding value estimation for the selection of cows to be kept in the next cycle (“pedigree_BVE”)
2. Genotyping of the calves and subsequent selection via single step BLUP on the farm (“ssBLUP_BVE”)
3. Reducing the selection intensity in the breeding company by 50% (“Low_SelectionIntensity”)
4. Modification of the selection index to put less weight on milk gain (RZM: 30, RZE: 20, RZR: 15, RZS: 10, RZKm: 5) (“Change_IndexWeights”)

To achieve reliable results, all scenarios were simulated 100 times and reported results represent averages across these runs. JSON-files of all scenarios can be found in Supplementary File S1. An overview of the development in the different scenario for the traits and the rates of inbreeding is given in Figure 6. By the use of a breeding value estimation (“pedigree_BVE” / “ss-BLUP_BVE”) all traits can be considered for selection, even when no phenotypes are available for the respective cohort. Even though both scenarios lead to a statistically significant increase in RZM (t-test, p = 0.0096 / 0.033, Figure 6.A) the practical differences between scenarios are small, as most of the genomic gain is obtained via the male side of the breeding program. Obtained prediction accuracies for both prediction methods are relatively low (e.g. for RZM around 0.42 for calves). To increase the effect of such selection, one could further consider selecting the cows used for reproduction as only half of the kept cows are used for mating to generate new animals for the calf+ cohort. For this, however, it would be required to use sexed semen.

**Figure 6.**
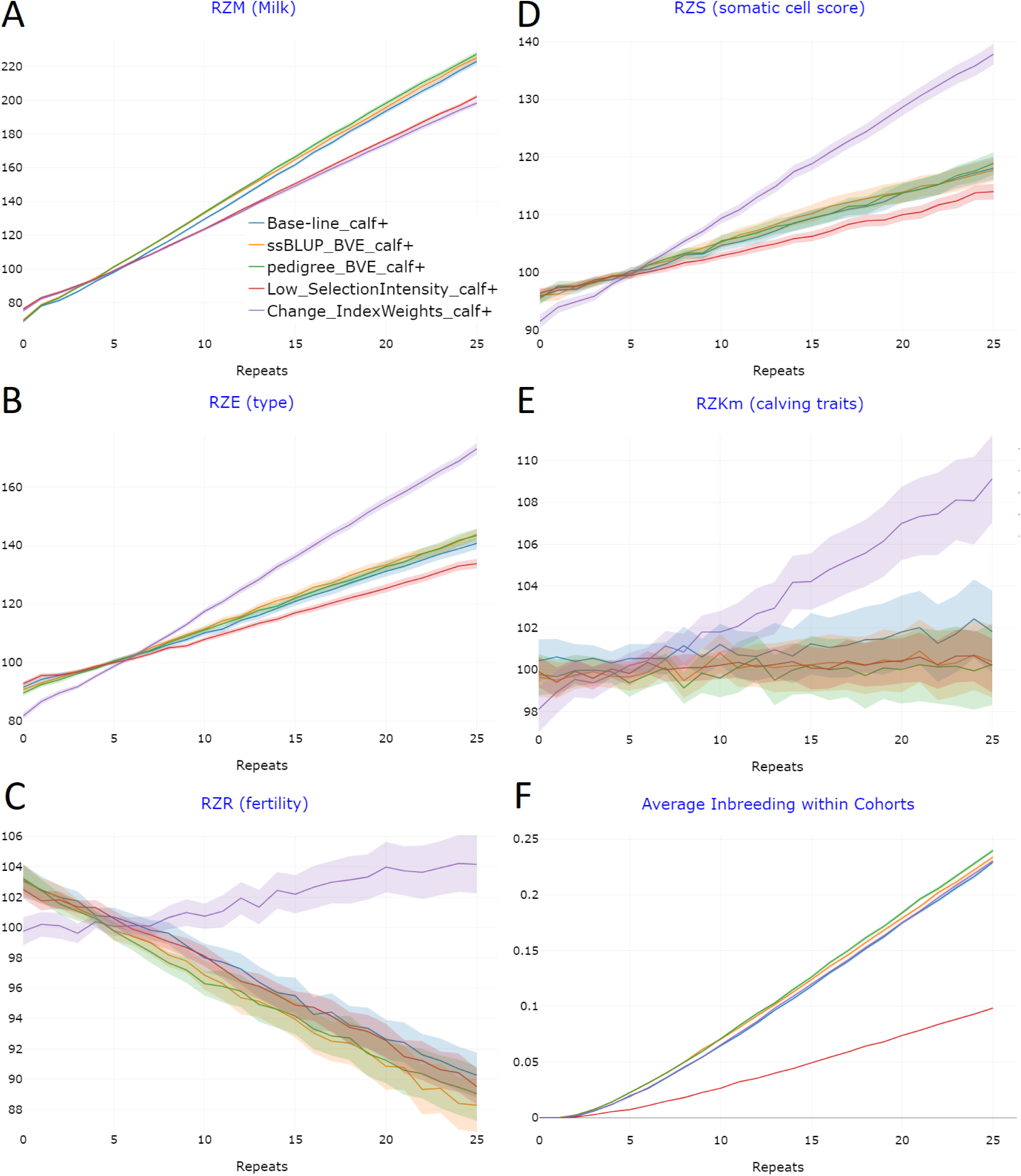
Genomic gain and the increase in inbreeding for the different scenarios of the cattle breeding program with 95% confidence bands for the traits RZM (A), RZE (B), RZR (C), RZS (D), RZKm (E), and inbreeding rates (F). Genomic values for all traits were standardized to an average of 100 after 5 repeats. This figure is an exemplary output of the “CompareProject”-module in MoBPSweb (www.mobps.de).

The reduction of the selection intensity on the side of the breeding company by 50% is leading to a reduction of the inbreeding rates by 57%, resulting in inbreeding rates of less than 0.005 per repeat (Figure 6.F). This however comes at the cost of a 20% reduction of genetic progress for RZM (Table 3, Figure 6.A). However, the ratio between genomic gain and inbreeding is highest in this scenario. Note that when considering a longer time horizon, genetic progress in this scenario might be highest as genomic gains in other scenarios will potentially drop due to lower remaining genetic diversity. To analyze this in detail, a higher number of repeats would have to be simulated. Furthermore, potentially counter measurements like the introduction of new diversity into the breeding nucleus should be considered then.

**Table 3.** Average genomic values and rates of inbreeding after 25 cycles of breeding for the newly generated calves with genomic values being standardized to 100 after 5 cycles. Numbers in brackets indicate the estimated standard deviation of the obtained averages.

| Scenario | RZM | RZE | RZR | RZS | RZKm | inbreeding |
| --- | --- | --- | --- | --- | --- | --- |
| Base-line | 223.0 (0.97) | 140.7 (1.05) | 90.3 (0.76) | 118.1 (1.00) | 101.8 (1.00) | 0.229 (0.00082) |
| pedigree_BVE | 227.3 (0.85) | 143.3 (1.15) | 89.0 (0.91) | 118.9 (0.98) | 100.3 (0.99) | 0.239 (0.00087) |
| ssBLUP_BVE | 226.1 (0.89) | 142.4 (0.95) | 88.5 (0.90) | 119.1 (1.01) | 100.7 (0.88) | 0.234 (0.00090) |
| Low_SelectionIntensity | 202.1 (0.62) | 133.8 (0.86) | 89.5 (0.61) | 114.0 (0.71) | 100.2 (0.66) | 0.099 (0.00035) |
| Change_IndexWeights | 198.3 (0.98) | 173.0 (1.50) | 104.2 (0.48) | 137.9 (0.85) | 109.1 (0.29) | 0.230 (0.00079) |

Modification of the selection index to put lower weight on the RZM is leading to a 22% smaller increase for this trait (Table 3). However, for all other traits performance is best, as this is the only scenario where an improvement for RZR was obtained and genomic gains for RZS were doubled (Figure 6.C/D).

Computing times for each simulation were about 20.5 minutes with about 15 minutes used for breeding value estimation, 4 minutes to generate new animals, and 30 seconds for both JSON-to-R conversion and initialisation of the founder population using a single core of an Intel(R) Xeon(R) Gold 6132 CPU 2.60 GHz.

## Discussion

MoBPSweb is providing an interactive, flexible and efficient platform to simulate, evaluate and compare breeding programs. Simulation studies are a valuable tool for breeders to quantify effects of their chosen breeding actions. Even though expectations regarding the direct effect of a change in the breeding program can usually be given based on experience and selection theory, breeding programs by nature are highly complex and breeding actions are interconnected, thereby affecting other parts of the breeding program. By the use of a simulation study a variety of output variables of a breeding program can be analyzed jointly, allowing to select the best solution for achieving a given breeding objective and thus optimize the breeding program. This solutions will of course be highly dependent on the breeding objective, the general framework and potential auxiliary conditions.

More fundamentally, the concepts introduced in Simianer *et al*. (2020) and the input environment given via MoBPSweb are providing a standardized, unambiguous and reproducible way to describe breeding programs and by this is solving common problems of unclear terminology for breeding programs. Thereby, the described framework can also be seen as a management tool for breeding programs in general and be used for planning costs, time and resources required to implement the program.

MoBPSweb is providing a variety of features of the MoBPS R-package (Pook *et al*. 2020). Note that a set environment with fixed modules and options to choose from does not offer the same degree of flexibility as the R-package. In particular, traits with complex phenotypes like longevity in cattle and sophisticated selected schemes that are using new and/or own methodology are usually not easy to include in MoBPSweb, so we still recommend using the MoBPS R-package in such cases.

Finally, note that no simulation study will be able to fully capture reality in its entirety. Nevertheless, the key strength of a simulation approach lies in the fact that, in contrast to field trials, far less time and money is needed to carry it out and potential harm to animals, such as adverse fitness effects, are avoided. Furthermore, experiments can be repeated and modified without any problems, which leads to much higher statistical power when comparing scenarios. Even if the absolute number of estimated effects might be off due to simplifications of reality, these effects should usually impact all scenarios and thereby still ensure comparability between scenarios.

## Acknowledgments

The MoBPS framework was developed in the context of the European Union’s Horizon 2020 Research and Innovation Program under grant agreement n°677353 IMAGE.

## Literature Cited

Cabrera-Bosquet, L., J. Crossa, J. von Zitzewitz, M. D. Serret, and J. Luis Araus, 2012 High-throughput Phenotyping and Genomic Selection. Journal of integrative plant biology 54: 312–320.

Chodorow, K., 2013 MongoDB: The definitive guide: powerful and scalable data storage. O’Reilly Media, Inc..

Danecek, P., A. Auton, G. Abecasis, C. A. Albers, E. Banks, et al., 2011 The variant call format and VCFtools. Bioinformatics 27: 2156–2158.

Faux, A.-M., G. Gorjanc, R. C. Gaynor, M. Battagin, S. M. Edwards, et al., 2016 AlphaSim: Software for breeding program simulation. The Plant Genome 9: doi: 10.3835/plantgenome2016.02.0013.

Fisher, R. A., 1918 XV. - The correlation between relatives on the supposition of Mendelian inheritance. Earth and Environmental Science Transactions of the Royal Society of Edinburgh 52: 399–433.

Galton, F., 1889 Natural Inheritance. Macmillan and Company.

Henryon, M., P. Berg, and A. C. Sørensen, 2014 Animal-breeding schemes using genomic information need breeding plans designed to maximise long-term genetic gains. Livestock Science 166: 38–47.

Jinek, M., K. Chylinski, I. Fonfara, M. Hauer, J. A. Doudna, et al., 2012 A programmable dual-RNA–guided DNA endonuclease in adaptive bacterial immunity. Science 337: 816–821.

Klein, R. J., C. Zeiss, E. Y. Chew, J.-Y. Tsai, R. S. Sackler, et al., 2005 Complement factor H polymorphism in age-related macular degeneration. Science 308: 385–389.

Legarra, A., O. F. Christensen, I. Aguilar, and I. Misztal, 2014 Single Step, a general approach for genomic selection. Livestock Science 166: 54–65.

Liu, H., B. B. Tessema, J. Jensen, F. Cericola, J. R. Andersen, et al., 2019 ADAM-plant: A software for stochastic simulations of plant breeding from molecular to phenotypic level and from simple selection to complex speed breeding programs. Frontiers in plant science 9: 1926.

Mendel, G., 1866 Versuche uber Pflanzen-Hybriden. Verhandlungen des Naturforschenden Vereines in Brünn 4: 3–47.

Meuwissen, T. H. E., B. J. Hayes, and M. E. Goddard, 2001 Prediction of total genetic value using genome-wide dense marker maps. Genetics 157: 1819–1829.

Oyama, K., T. Katsuta, K. Anada, and F. Mukai, 2002 Heritability and repeatability estimates for reproductive traits of Japanese Black cows. Asian-australasian journal of animal sciences 15: 1680–1685.

Pérez-Enciso, M., L. C. Ramírez-Ayala, and L. M. Zingaretti, 2020 SeqBreed: A python tool to evaluate genomic prediction in 67 complex scenarios. Genetics Selection Evolution 52: 1–9.

Pook, T., M. Schlather, and H. Simianer, 2020 MoBPS - Modular Breeding Program Simulator. G3: Genes, Genomes, Genetics 10: 1915–1918.

Purcell, S., B. Neale, K. Todd-Brown, L. Thomas, M. A. R. Ferreira, et al., 2007 PLINK: A tool set for whole-genome association and population-based linkage analyses. The American Journal of Human Genetics 81: 559–575.

R Core Team, 2017 R: A Language and Environment for Statistical Computing.

Roman, R. M., C. J. Wilcox, and F. G. Martin, 2000 Estimates of repeatability and heritability of productive and reproductive traits in a herd of Jersey cattle. Genetics and Molecular Biology 23: 113–119.

Sargolzaei, M. and F. S. Schenkel, 2009 QMSim: A large-scale genome simulator for livestock. Bioinformatics 25: 680–681.

Sidnell, P., 2007 Warhorse: Cavalry in ancient warfare. Bloomsbury Publishing.

Simianer, H., A. Ganesan, L. Büttgen, N.-T. Ha, and T. Pook, 2020 A unifying concept of animal breeding programs. Companion Paper.

Solberg, L. C., W. Valdar, D. Gauguier, G. Nunez, A. Taylor, et al., 2006 A protocol for high-throughput phenotyping, suitable for quantitative trait analysis in mice. Mammalian Genome 17: 129–146.

VanRaden, P. M., 2008 Efficient methods to compute genomic predictions. Journal of Dairy Science 91: 4414–4423.

Vereinigte Informationssysteme Tierhaltung w.V., 2020 Estimation of Breeding Values for Milk Production Traits, Somatic Cell Score, Conformation, Productive Life and Reproduction Traits in German Dairy Cattle.

Wright, S., 1922 Coefficients of inbreeding and relationship. The American Naturalist 56: 330–338.

Zerbino, D. R., P. Achuthan, W. Akanni, M. R. Amode, D. Barrell, et al., 2018 Ensembl 2018. Nucleic acids research 46: D754–D761. 103

